# Low-intensity ultrasound drives offline neuromodulation through passive membrane mechanisms in synaptically isolated hippocampal neurons

**DOI:** 10.64898/2026.09.28.754934

**Authors:** Marcell Stippinger, Édua Édes, Barnabás Kovács, Aletta Mészáros, Veronika Házi, Melinda E. Gazdik, Zoltán Somogyvári, Attila Szűcs, Tamás Kiss

**Affiliations:** Theoretical Neuroscience and Complex Systems Research Group, Department of Computational Sciences, HUN-REN Wigner RCP, Budapest, Hungary; Doctoral College, Semmelweis University, Budapest, Hungary; Department of Physiology and Neurobiology, Eötvös Loránd University, Budapest, Hungary

**Keywords:** low-intensity focused ultrasound, ultrasound neuromodulation, offline effects, membrane noise, membrane capacitance, leak reversal potential, membrane time constant, hippocampal cell culture, patch-clamp, intrinsic excitability

## Abstract

Low-intensity ultrasound alters neuronal excitability long after exposure ends, but the basis of this offline effect is unclear and most proposed mechanisms require synaptic receptors. We recorded from cultured mouse hippocampal neurons in whole-cell current clamp, before and 20 min after 1 min of 850 kHz continuous-wave ultrasound at 0.5 W/cm^2^ or sham, with synaptic transmission blocked throughout, each neuron re-patched. Sonicated neurons remained intact and firing, yet depolarized, with smaller action potentials and more noise. A passive-membrane fit showed reduced capacitance and a depolarized leak reversal potential in every sonicated cell, with a shortened time constant and no detectable change in input conductance, while sham cells showed no significant change. Changes in the time constant and reversal potential, both independent of membrane area, indicate a modification of the membrane itself rather than a loss of area. Brief ultrasound therefore leaves a lasting, receptor-independent imprint on passive membrane properties.

## 1. Introduction

The history of ultrasound neuromodulation can be traced back to early 20^th^-century observations of the biological effects of acoustic energy on cellular tissues (Harvey and Loomis, 1928). However, the foundational principles of the field were established in the mid-1950s, when Fry and colleagues systematically demonstrated that focused ultrasound could reversibly modulate neural activity within the central nervous system (Fry, 1953; Fry et al., 1958). After decades of intermittent progress, the field has undergone a major resurgence driven by the refinement of low-intensity focused ultrasound (LIFU) (Tufail et al., 2010; Baek et al., 2017; Fomenko et al., 2018). This modern approach offers a non-invasive, high-resolution method for neuromodulation, shifting the paradigm from irreversible surgical ablation toward the precise, reversible manipulation of neural circuits (Blackmore et al., 2019; Osada and Konishi, 2024). Today, LIFU is regarded as a promising tool in both basic neuroscience and clinical neurology, supported by rapidly expanding translational interest (Darmani et al., 2022; Collins and Mesce, 2022).

Recent years have seen a proliferation of studies across diverse paradigms, ranging from *in vitro* cell cultures to transcranial human applications (Shi et al., 2025). Current work addresses questions from parameter optimization to potential clinical use cases. Despite this progress, however, the fundamental mechanism of action of LIFU remains debated, particularly with respect to the factors that distinguish immediate, transient effects from long-lasting cellular changes. Although it is generally accepted that LIFU may act through mechanical, thermal, or cavitational effects (Dell’Italia et al., 2022), how these physical influences are translated into multiscale changes spanning subcellular, cellular, and network levels remains poorly understood.

The field of transcranial ultrasound stimulation (TUS) has recently shifted its focus from transient “online” effects to the induction of sustained, “offline” neuroplasticity. This transition is driven by the therapeutic promise of TUS to produce long-lasting circuit modulation in neurological and psychiatric disorders. For example, in humans, Zeng et al. (2022) demonstrated that an 80-second train of theta-burst TUS (tbTUS) applied to the motor cortex induced a significant increase in corticospinal excitability that persisted for at least 30 to 60 minutes. This evidence for “long-term potentiation-like” plasticity was recently expanded by findings showing that TUS can also bidirectionally modulate brain function. By mimicking endogenous brainwave patterns such as theta and gamma oscillations, Kim et al. (2024) showed that TUS can entrain neural activity to induce either long-term potentiation- or long-term depression-like shifts in excitability that persist well beyond the stimulation period.

While these systems-level studies underscore the clinical promise of TUS, they also highlight the need to identify the cellular and molecular mechanisms underlying sustained effects. To gain a more mechanistic understanding, Kim et al. (2017) used organotypic rat hippocampal slice cultures and showed that spike counts increased specifically after, rather than during, ultrasound application relative to sham stimulation. This “offline” modulation was further characterized by Clennell et al. (2021), who demonstrated in primary rat cortical neurons that a single 40 s ultrasound exposure enhanced excitability for up to 8 h. Building on this, Clennell et al. (2023) identified a specific molecular mechanism for these shifts, showing that ultrasound induces sustained increases in whole-cell potassium currents through signaling cascades triggered by ionotropic glutamate receptors. Similarly, Fan et al. (2024) found in cultured hippocampal neurons that low-intensity pulsed ultrasound (LIPUS) significantly increased spontaneous action-potential frequency and excitatory synaptic currents (sEPSCs) for more than 10 minutes. They reported that this effect is mediated by facilitation of L-type calcium channels, which subsequently activates the CaMKII–CREB pathway to regulate long-term gene transcription. While these studies focus on excitatory effects, recent evidence from Mishima et al. (2024) highlights a divergent, persistent inhibitory response: repetitive pulsed-wave ultrasound suppressed spontaneous recurrent burst firing in hippocampal cultures for several minutes after the stimulation period ended. This effect was mediated by astrocytic TRPA1 activation and a subsequent increase in ambient GABA levels, demonstrating that the offline state can also be shaped by non-neuronal cells within the network. Furthermore, Niu et al. (2022) showed that 5 minutes of transcranial focused ultrasound targeting the hippocampal perforant path induced a sustained depression of synaptic transmission that persisted for at least 30 minutes, with effect magnitude depending on pulse repetition frequency. This circuit-level picture was recently extended by Watts et al. (2025), who found that brief theta-burst ultrasound causes lasting modifications to hippocampal synaptic circuitry and plasticity susceptibility via a canonical Akt-dependent metaplastic process. Direct measurements made during sonication are rarer. In isolated crayfish axon, Lin et al. (2019) recorded an ultrasound-gated conductance with a directly measured reversal potential under synaptic blockade, an effect that rose and decayed with the stimulus rather than outlasting it. Together, these findings suggest that sustained ultrasound effects arise from a complex interplay among intrinsic neuronal properties, synaptic signaling, and glial interactions.

These divergent findings—ranging from sustained synaptic depression to prolonged cellular hyper-excitability and astrocyte-mediated inhibition—highlight both the complexity of the phenomenon and a fundamental ambiguity in the field concerning the offline effects of ultrasound treatment. To reduce this complexity and isolate neuronal components, we built on the methodological framework of Clennell et al. (2021) and examined synaptically isolated individual neurons in mouse hippocampal cultures. This approach allowed us to bypass network-level interactions and investigate whether sustained changes in excitability after ultrasonic stimulation can be explained by cell-intrinsic membrane mechanisms.

## 2. Methods

### 2.1. Ethical approval

Experiments were carried out in accordance with the Hungarian Act of Animal Care and Experimentation (1998, XXVIII), Directive 2010/63/EU of the European Parliament and of the Council of 22 September 2010 on the protection of animals used for scientific purposes, and the European Convention for the Protection of Vertebrate Animals Used for Experimental and Other Scientific Purposes. Under this legislation, killing animals to obtain brain tissue for primary neuronal cultures is regarded as organ donation, so no individual ethical approval was required for this study. Animal husbandry and handling at the host laboratory are authorized by the institutional Ethical Committee on Animal Experimentation and the National Scientific Ethical Committee on Animal Experimentation (authorization #PEI/001/1108-4/2013). All efforts were made to minimize the number of animals used and their suffering.

### 2.2. Animals and primary hippocampal neuronal cultures

CD-1 wild-type mice (CD-1^®^ IGS Mouse, strain code 022; Charles River Laboratories, Wilmington, MA, USA) were housed at 22 ± 1 °C under a 12-hour light/dark cycle with food and water available *ad libitum*. For breeding, adult males and females were paired and co-housed for four days, during which the females were checked daily for a vaginal plug to confirm mating. Pregnant dams were monitored daily until tissue collection.

Primary hippocampal cultures were prepared from CD-1 embryos at embryonic day 16–19 (E16– 19); the sex of the embryos was not determined. Pregnant animals were killed by rapid cervical dislocation without anesthesia. The embryos were decapitated, and the brains were rapidly removed and the hippocampi dissected under aseptic conditions, following established protocols (Czöndör et al., 2009; Oueslati Morales et al., 2021). The hippocampal tissue was digested with trypsin-EDTA and DNase I in phosphate-buffered saline (PBS; Gibco, Thermo Fisher Scientific) for 15 min at 37 °C. Digestion was stopped by adding heat-inactivated fetal calf serum (FCS), followed by centrifugation at 400 × *g* for 10 min. The pellet was dissociated by gentle trituration, and the suspension was filtered through a gamma-sterilized polystyrene mesh (42 µm pore size; EmTek, Hungary). Viable cells were counted with trypan blue in a Bürker chamber.

Cells were seeded at 7 × 10^4^ cells*/*cm^2^ onto plasma-cleaned (Zepto plasma cleaner, Diener; 100 W, 10 min), round glass coverslips (12 mm diameter) placed in 24-well plates (CELLSTAR, Greiner Bio-One) and coated with poly-L-lysine and laminin (both Sigma-Aldrich). The starting medium was Neurobasal Plus (cat. no. A3582901; Thermo Fisher Scientific, Waltham, MA, USA) supplemented with 2 % B27 Plus (cat. no. A3582801), 5 % FCS (PAN Biotech, Aidenbach, Germany) and 0.5 mM GlutaMAX (cat. no. 35050038; Thermo Fisher Scientific). All cultures were maintained at 37 °C in a humidified atmosphere containing 5 % CO_2_. On day in vitro 1 (DIV1) the medium was completely replaced with fresh starting medium, and 10 µM cytosine *β* -D-arabinofuranoside (Sigma-Aldrich) was added between DIV3 and DIV5 to inhibit glial cell division. On DIV5, one third of the medium was replaced with BrainPhys (cat. no. 05790; StemCell Technologies, Vancouver, BC, Canada) supplemented with 2 % NeuroCult SM1 (cat. no. 05711; StemCell Technologies), and on DIV9 and DIV12 half of the conditioned medium was replaced with the same fresh medium. Electrophysiological recordings were performed at DIV10–12, allowing sufficient time for neurons to acquire mature electrophysiological characteristics. Recordings were collected on three experimental days, each using a separate culture preparation: the eight sonicated neurons came from two preparations (four cells each) and the eight sham neurons from a third, so that cells sharing a treatment group and a recording day also shared a dish.

### 2.3. Electrophysiological recordings and ultrasound stimulation

Whole-cell patch-clamp experiments (*n*_US_ = 8 ultrasound-sonicated neurons and *n*_S_ = 8 sham-sonicated neurons) were performed at room temperature (22–24 °C) under an Axiovert 200 inverted microscope (Zeiss) with differential interference contrast (DIC) optics. Evoked responses were recorded in whole-cell current-clamp configuration using a MultiClamp 700B amplifier (Molecular Devices).

Intracellular voltage and current signals were digitized at 20 kHz. Stimulus command waveforms were generated using DASYLab v11 (DASYTec USA, National Instruments). Patch pipettes were pulled from borosilicate glass capillaries and had resistances of 7–10 MΩ when filled with intracellular solution.

Access resistance was compensated by bridge balancing before recordings. Throughout the experiment, cultures were bathed in artificial cerebrospinal fluid (ACSF) containing pharmacological blockers of synaptic transmission to isolate intrinsic membrane properties.

The bath solution contained (in mM): NaCl 140, KCl 5, CaCl_2_ 2, MgCl_2_ 1, HEPES 5, and D-glucose 10; pH was adjusted to 7.45. Patch electrodes were filled with the following solution (in mM): K-gluconate 100, KCl 10, KOH 20, MgCl_2_ 2, NaCl 2, HEPES 10, EGTA 0.2, and D-glucose 5; pH was adjusted to 7.3. Neurons were synaptically isolated using the following receptor blockers: CNQX (10 µM), AP5 (40 µM), and bicuculline (30 µM). All three blockers were present continuously from before the first recording until the end of the second, in the recording chamber and in the dish used for sonication, so receptor antagonism was maintained during the transfer and during the ultrasound exposure itself.

To characterize intrinsic membrane properties, neurons were subjected to stepwise current injections from − 200 to 200 pA in 5 pA increments, with each stimulus cycle lasting 1.25 s. Voltage and current responses were stored for offline analysis.

Following the initial whole-cell recording, the coverslip containing the primary culture was transferred to a Petri dish filled with the same blocker-containing ACSF for sonication at room temperature, 22–24 °C. Ultrasound was generated with an in-house-built signal generator, amplifier, and transducer (Appendix, Sec. A1.1) and applied for 1 min as a continuous wave at 850 kHz with a peak intensity of *I*_SPPA_ = 0.5 W*/*cm^2^. The ultrasound pressure field (Fig. 1C) was characterized using a needle hydrophone

**Figure 1.**
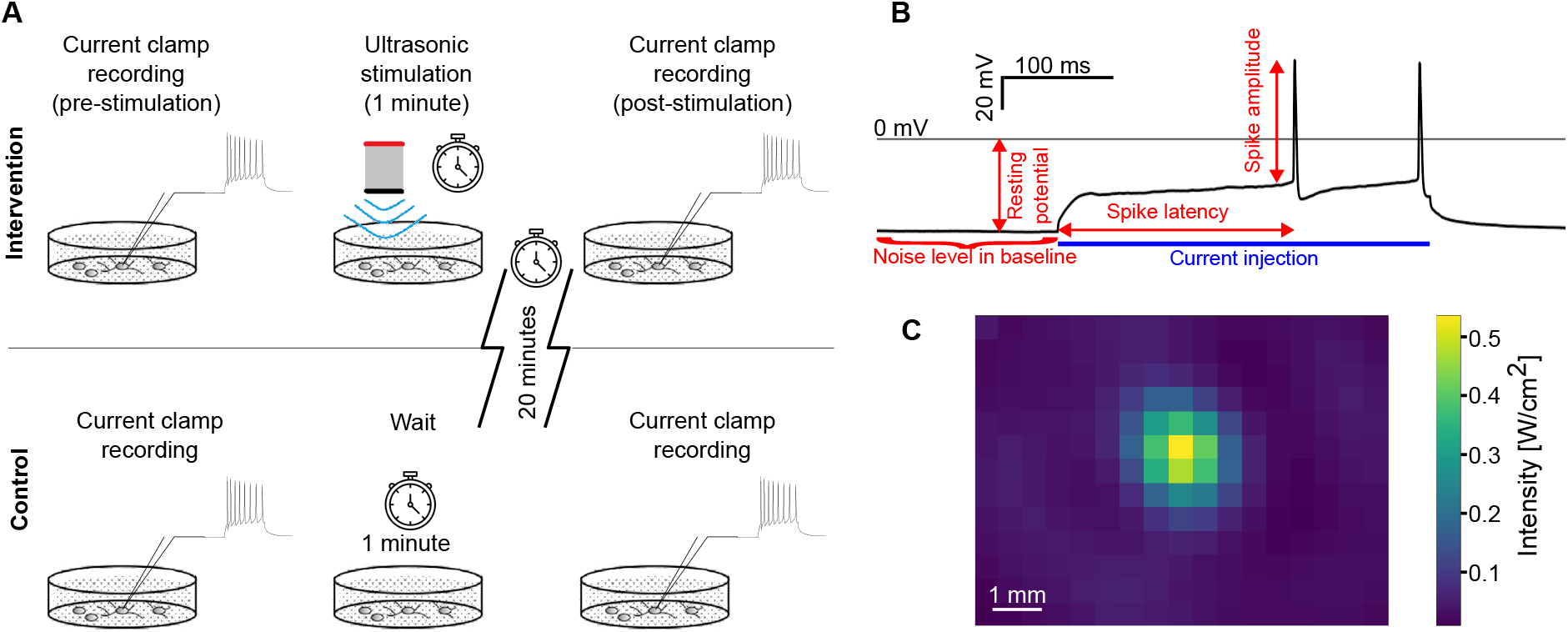
*In vitro* experimental setup and analyzed measures. A. Experimental protocol showing ultrasound (intervention) and sham (control) conditions. Whole-cell current-clamp recordings were performed in the same neurons before and approximately 20 min after a 1 min-long sonication session. B. Membrane-potential characteristics were calculated from baseline voltage recordings and responses to positive current injection. C. Ultrasound intensity measured at the depth of the coverslip with the cell culture (5 mm below the surface).

(HNR-0500 (74145 V1), ONDA, Sunnyvale, CA, USA) and an in-house-built spatial scanner. The coverslip was then returned to the recording chamber, previously patched neurons were re-identified, and a second whole-cell recording was performed. Transfer, sonication, and return required approximately 20 minutes in total (Fig. 1A).

### 2.4. Passive membrane parameter estimation

Passive membrane properties were estimated from the responses to hyperpolarizing current steps, where voltage-gated conductances are minimally engaged and the membrane behaves as a passive RC circuit,

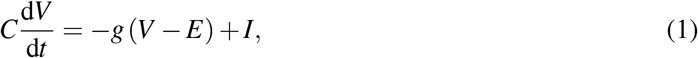

whose response to a current step is a single exponential relaxation,

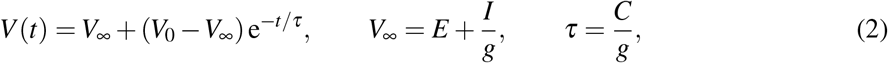

where *g* is the input (leak) conductance, *E* the leak reversal potential, *C* the membrane capacitance, *τ* the membrane time constant, *V*_∞_ the steady-state voltage approached during the step, and *V*_0_ = *V* (*t* = 0) the voltage at step onset.

For each hyperpolarizing sweep (current steps between −210 pA and −50 pA), the membrane potential over the 0.5–50 ms window following step onset was fitted to this exponential by nonlinear least squares (`scipy.optimize.curve_fit`), with *V*_0_ fixed to the measured voltage at the window start and *V*_∞_ and *τ* as free parameters. For each cell and condition, the steady-state voltages *V*_∞_ were then regressed on the injected current *I* by ordinary least squares: the input conductance was obtained as the inverse of the slope, *g* = (d*V*_∞_*/*d*I*)^−1^, and the reversal potential *E* as the intercept. The membrane capacitance was computed as 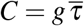, where 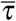 is the mean of the per-sweep time constants *τ* for that cell and condition. Estimating *g* and *E* from the current–voltage relationship and *C* from the membrane time constant identifies the three parameters from distinct features of the recording rather than from a single trace. Throughout, the recording pipette was positioned on the soma, the reference electrode was placed in the same bath but away from the recorded neuron, and the recording dish rested on a grounded metal stage. The same current steps, fitting window and estimator were applied to a given neuron before and after treatment.

### 2.5. Data analysis and statistics

Voltage responses to current-step stimulation were used to extract multiple physiological parameters, including resting membrane potential (RMP), action-potential (AP) latency, and AP amplitude, using custom software (Szűcs, 2022) (Fig. 1B). The AP threshold was defined as the voltage level, at which the membrane potential slope exceeded 5 mV*/*ms; AP amplitude was then calculated as the voltage difference between threshold and peak. For analyses of repetitive firing during positive current steps, the first evoked spike was used for measurement and values were averaged across positive steps. To characterize latency—that is, the time lag between the onset of the current step (*I*) and the first action potential—we fitted the first-spike latencies with a Bělehrádek function (Bělehrádek, 1935):

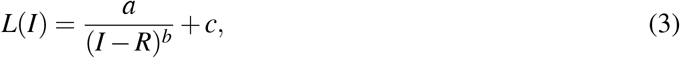

where *R* is the rheobase current (the current eliciting the first action potential with infinite latency) and *a, b*, and *c* determine the scale, power-law decay, and asymptotic latency of the curve. For standardized comparison across cells, latency was evaluated from the fitted curve at 1.3 times the observed rheobase. RMP was the mean membrane potential over the 2 ms preceding each current step.

Membrane-potential noise was measured from the unstimulated baseline preceding each current step, where no current was injected. For every step we took the − 600– 50 ms window relative to step onset, which excludes both the relaxation transient left by the preceding step and the onset of the next one, removed a second-order polynomial trend, and computed the standard deviation of the residual. Windows containing a residual action potential (peak membrane potential above − 20 mV) were discarded. The value reported for a recording is the median across its steps (57–91 steps per recording).

Only neurons that were successfully re-patched after treatment were analyzed; of 21 neurons recorded before treatment (ultrasound and sham together), 5 could not be re-patched and were excluded. Cells with an unstable baseline or a lack of giga-seal at the first patch were also excluded from the analysis. Coverslips were assigned randomly to ultrasound or sham treatment by É.É.; the analysis was not performed blind to treatment.

We used statistical procedures appropriate for small samples and non-normal data. Within each treatment group (sonicated and sham), before–after changes were tested with paired exact Wilcoxon signed-rank tests and exact sign tests on paired differences. To compare treatment effects between groups, we calculated per-cell treatment deltas for each measured feature and compared the resulting distributions using exact permutation tests on median delta values and Mann–Whitney U tests. Effect sizes were quantified using Cliff’s delta, a distribution-free measure suitable for small samples. All *p*-values were corrected for multiple testing within each family of measured features using the Benjamini–Hochberg false discovery rate (FDR) procedure with *α* = 0.05.

Spike number in response to injected current (I–F) was analyzed with linear mixed-effects models and repeated-measures ANOVA. Spike number was modeled as a function of injected current (continuous covariate), condition (before vs. after treatment), and their interaction, so that the current×condition term tested directly whether treatment altered input–output gain. Separate models were fit for the ultrasound and sham groups, followed by a combined model including all main effects and pairwise interactions of treatment group, condition, and current to assess whether treatment-specific effects differed between groups; findings were verified with a repeated-measures ANOVA using individual cells as a blocking factor. Analyses included all measurements with injected current from 20 pA, up to the highest current shared across all 32 experiments (8 cells × 2 treatments × 2 conditions), with reference levels “before” and “sham” and significance assessed at *α* = 0.05. All models were fit using the `statsmodels` Python library.

The use of generative AI tools in preparing this work is described in the Generative AI Statement.

## 3. Results

### 3.1. Ultrasound-induced membrane parameter changes

To evaluate the impact of LIFU on the electrophysiological properties of primary mouse hippocampal neurons, individual cells were characterized before and after ultrasound stimulation or sham treatment. Morphological assessment by light microscopy revealed no discernible structural alterations or signs of cellular distress, such as blebbing or swelling, following repeated whole-cell recordings or ultrasound exposure (Fig. 2A). Functional viability was likewise preserved: neurons re-patched after ultrasound application retained stable resting membrane potentials and characteristic firing patterns consistent with healthy cells (Fig. 2B). Nevertheless, qualitative inspection of spontaneous voltage recordings suggested a subtle increase in baseline noise, prompting a more detailed quantitative analysis of membrane and firing properties.

**Figure 2.**
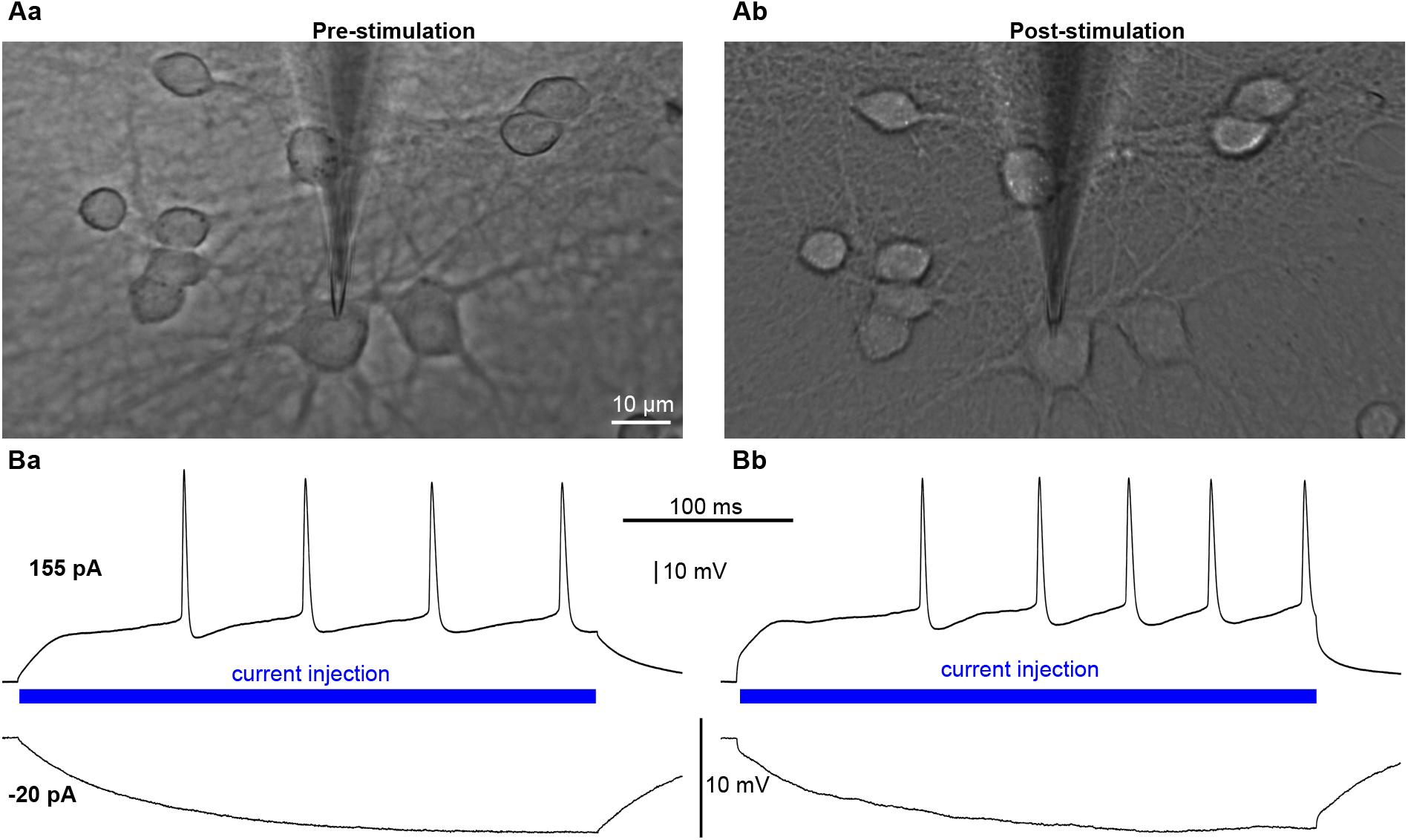
Morphological and functional stability of hippocampal neurons following LIFU. A. Representative light-microscopy images of a primary mouse hippocampal neuron before (Aa) and after (Ab) ultrasound stimulation, taken in differential interference contrast (DIC) on the recording microscope (Axiovert 200, Zeiss). No discernible structural alterations, such as membrane blebbing or somatic swelling, were observed following the ultrasound protocol and subsequent re-patching of the same cell. B. Electrophysiological signatures confirming functional viability. Current-clamp recordings show characteristic voltage responses to depolarizing (155 pA) and hyperpolarizing (− 20 pA) current steps before stimulation (Ba) and after stimulation (Bb). Despite a subtle increase in spontaneous baseline fluctuations after stimulation, neurons maintained robust action-potential firing and stable overall membrane behavior. Scale bars: 10 µm in Aa (Ab shows the same field at the same magnification); 10 mV and 100 ms in B.

LIFU produced consistent and significant changes in multiple electrophysiological properties, whereas sham-treated control cells showed no significant pre–post changes in any of the measured parameters (Fig. 3A–D). Within the sonicated group, RMP shifted significantly toward depolarization after treatment (median before: −56.8 mV, median after: −51.9 mV, median change: +5.9 mV; Wilcoxon signed-rank test, FDR-corrected *p* = 0.021, *n* = 8 pairs). Spike amplitude also decreased significantly after sonication (median before: 69.1 mV, median after: 59.6 mV, median change: − 8.0 mV; *p* = 0.021). Membrane noise increased markedly following sonication (median before: 91.9 µV, median after: 240.6 µV, median change: +153.8 µV, a 2.6-fold rise; Wilcoxon signed-rank test, *p* = 0.021). The change in spike latency in sonicated cells (median change: − 28.2 ms) was not significant after correction for multiple comparisons (*p* = 0.219). No significant pre–post changes were observed in any parameter in sham-treated cells (all *p* ≥ 0.738).

**Figure 3.**
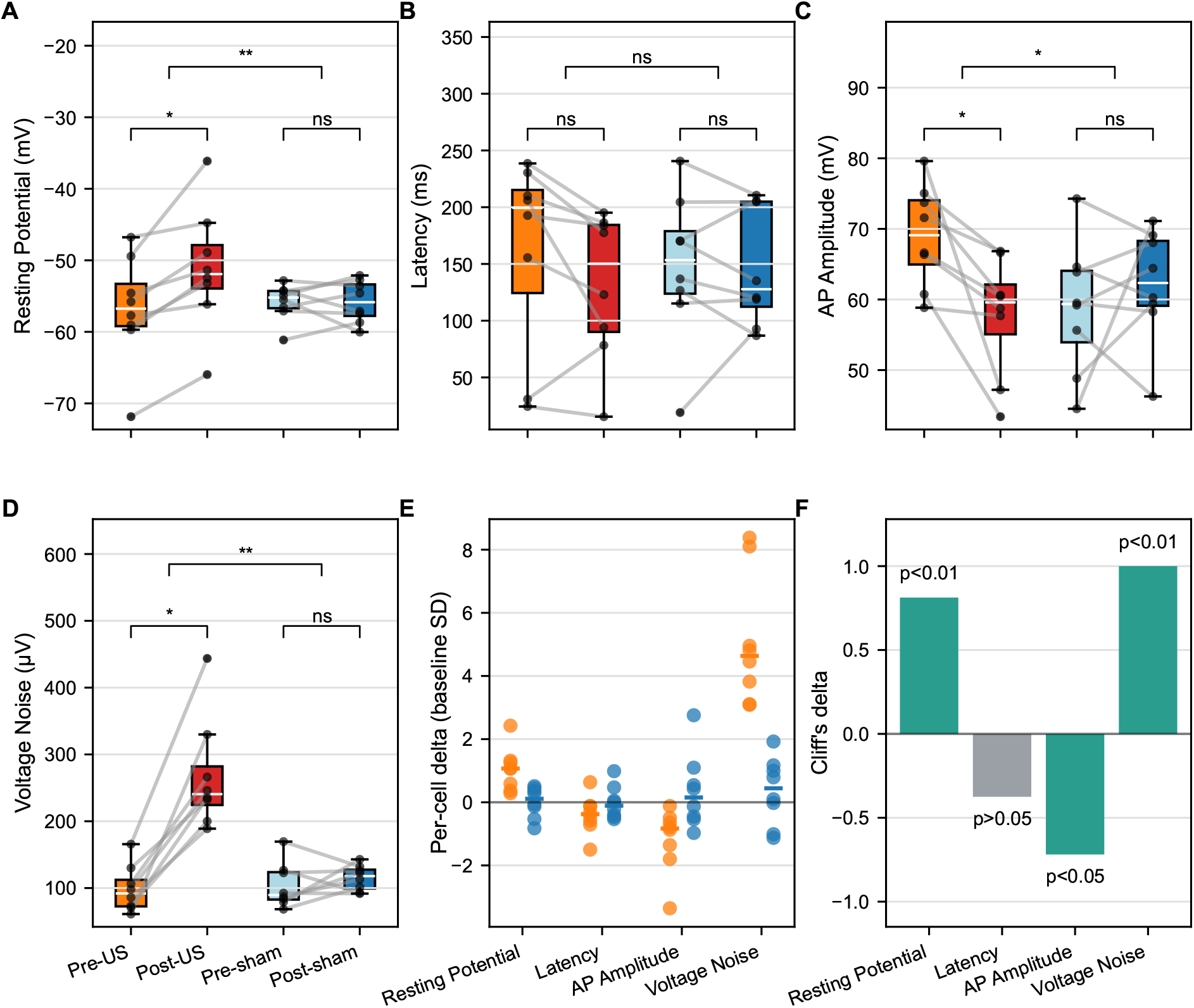
Effect of ultrasound stimulation on neuronal membrane parameters. A–D. Resting membrane potential, first-spike latency, spike amplitude, and membrane-potential noise during unstimulated baseline periods, respectively (∗ : *p* ≤ 0.05, ∗∗: *p*≤ 0.01; see Methods for details of metric extraction). Connected points represent individual neurons (*n* = 8) before and after sonication. Boxplots show the median, 25^th^ and 75^th^ percentiles, range, and outliers (points beyond 1.5 times the interquartile range). E. Comparison of treatment-induced changes (orange: ultrasound delta; blue: sham delta), each parameter divided by the standard deviation of its pre-treatment values across all cells so that quantities in different units share one axis; horizontal bars mark group medians. F. Effect-size summary. Statistical analysis supports a significant increase in membrane noise and resting membrane potential, a significant decrease in spike amplitude, and no significant change in spike latency after ultrasound. Reported *p* values are FDR-corrected, and are shown in panel F as thresholds; exact values are given in the text.

Direct comparison of treatment-induced deltas between the sonicated and control groups confirmed these effects (Fig. 3E–F). Sonicated cells exhibited significantly larger depolarization shifts than control cells (exact permutation test on median delta difference: *p* = 0.009; Cliff’s *δ* = 0.81), significantly greater reductions in spike amplitude (*p* = 0.023; Cliff’s *δ* =− 0.72), and markedly greater increases in membrane noise relative to control cells (*p* = 0.009; Cliff’s *δ* = 1.00). The effect on latency did not differ significantly between groups (*p* = 0.332). The large effect-size values indicate strong and consistent sonication-induced effects with little overlap between groups. For membrane noise the separation was complete, every sonicated cell showing a larger increase than every sham cell.

Next, we analyzed spike-frequency changes in response to injected current (I–F) following ultrasound application (Fig. 4). Within-group mixed-effects models showed that in ultrasound-treated cells, the post-treatment change in I–F slope was significant (current:condition coefficient = 0.0600, *p* = 0.00141), whereas the intercept shift was not (coefficient = − 0.559, *p* = 0.585). In sham cells, the slope change was smaller and did not reach significance (coefficient = 0.0338, *p* = 0.119), and no intercept shift was detected (coefficient = − 0.435, *p* = 0.713). Although the slope increase in ultrasound-treated cells was approximately 1.78-fold larger than in sham cells, the direct test of this difference—the three-way current:condition:treatment interaction in the combined mixed-effects model—was not significant (coefficient = 0.0261, *p* = 0.363). A sensitivity model allowing random slopes for individual cells likewise found no significant effect for this interaction (coefficient = 0.0261, *p* = 0.213); by contrast, the treatment:condition term in the repeated-measures ANOVA verification did reach significance (*F* = 4.308, *p* = 0.0387). Taken together, these results indicate that only ultrasound-treated cells showed a significant post-treatment increase in I–F slope gain; the mixed-effects models were insufficient to establish a statistically significant treatment-specific difference, although the ANOVA treatment:condition term was significant.

**Figure 4.**
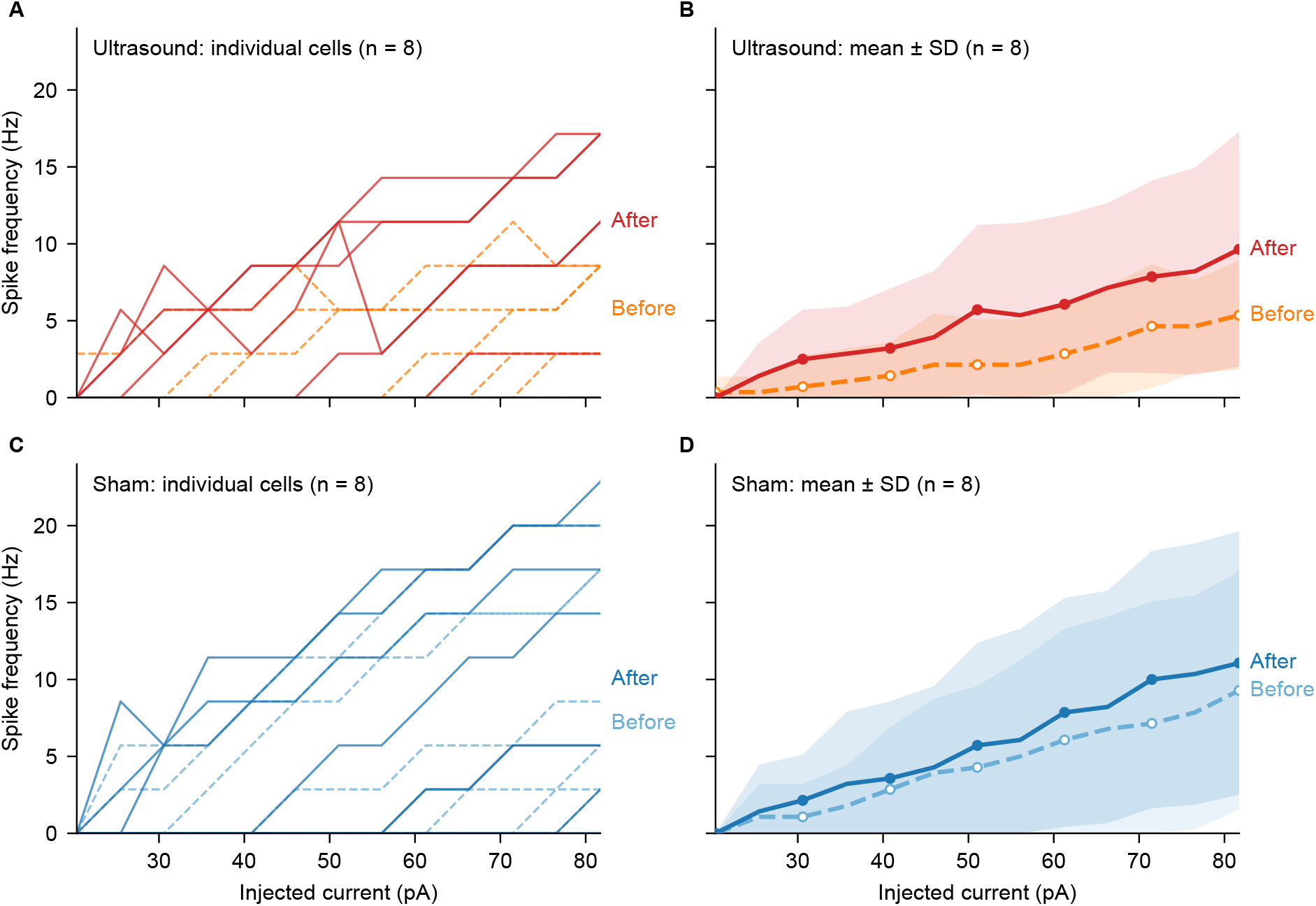
Spike frequency–current relationship before and after ultrasound (A, B) and sham treatment (C, D). A, C. Individual cells before (dashed lines) and after (solid lines) treatment; because spike counts per current step take discrete values, some curves coincide. B, D. Group means before (dashed lines, open circles) and after (solid lines, filled circles) treatment, with the shaded regions showing ± SD. Currents are restricted to the range recorded in every cell (20–82 pA). *n* = 8 cells per group, recorded before and after treatment: ultrasound, 8 cells from 2 culture preparations; sham, 8 cells from 1 culture preparation.

### 3.2. Passive membrane parameters

We estimated the passive membrane parameters—input conductance *g*, membrane capacitance *C*, and resting (leak) reversal potential *E*—from the exponential charging of the membrane during hyperpolarizing current steps (*n* = 8 cells per group; Fig. 5). Ultrasound produced no detectable change in the input conductance. Within the sonicated group *g* showed no significant paired change (median before 5.3 nS, after 5.4 nS; median Δ = +0.06 nS), and the between-group comparison was not significant (FDR-adjusted permutation *p* = 0.209; Cliff’s *δ* = − 0.56). The difference-in-differences was − 1.91 nS (95 % CI − 3.98 to 0.16 nS), an interval that includes zero, so a reduction of up to about 4 nS remains compatible with our measurements. Membrane capacitance, in contrast, decreased markedly after sonication (median before 139.6 pF, after 101.7 pF; median paired Δ = − 45.4 pF; all eight cells decreased, Wilcoxon *p* = 0.0078), whereas sham cells showed no significant change (median Δ = +7.3 pF; Wilcoxon *p* = 0.461); the treatment effect was significant between groups (FDR-adjusted permutation *p* = 0.0047; Cliff’s *δ* =−0.94). The reversal potential shifted in the depolarizing direction after sonication (median before − 63.1 mV, after − 56.5 mV; median paired Δ = +7.6 mV; all eight cells depolarized, Wilcoxon *p* = 0.0078), with no significant change in sham controls (median Δ = − 0.2 mV; Wilcoxon *p* = 0.641) and a significant between-group effect (FDR-adjusted permutation *p* = 0.0047; Cliff’s *δ* = +0.81). Together, these results indicate that ultrasound reduced membrane capacitance and produced a depolarizing shift in the resting reversal potential, without a detectable change in leak conductance. Because the capacitance is obtained as 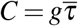 (see Methods), these two results can also be read directly in terms of the fitted time constant (Fig. 6). Within the sonicated group 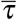 fell in seven of eight cells (median before 23.4 ms, after 18.0 ms; median Δ =−9.5 ms, a decrease of 31 %; Wilcoxon *p* = 0.016), whereas sham cells showed no change (median Δ = −0.5 ms; Wilcoxon *p* = 1.0). The between-group comparison of paired changes was not significant (permutation *p* = 0.056; Cliff’s *δ* = −0.59). We do not fold 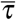 into the FDR family above, because 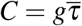 makes it algebraically dependent on parameters already tested there.

**Figure 5.**
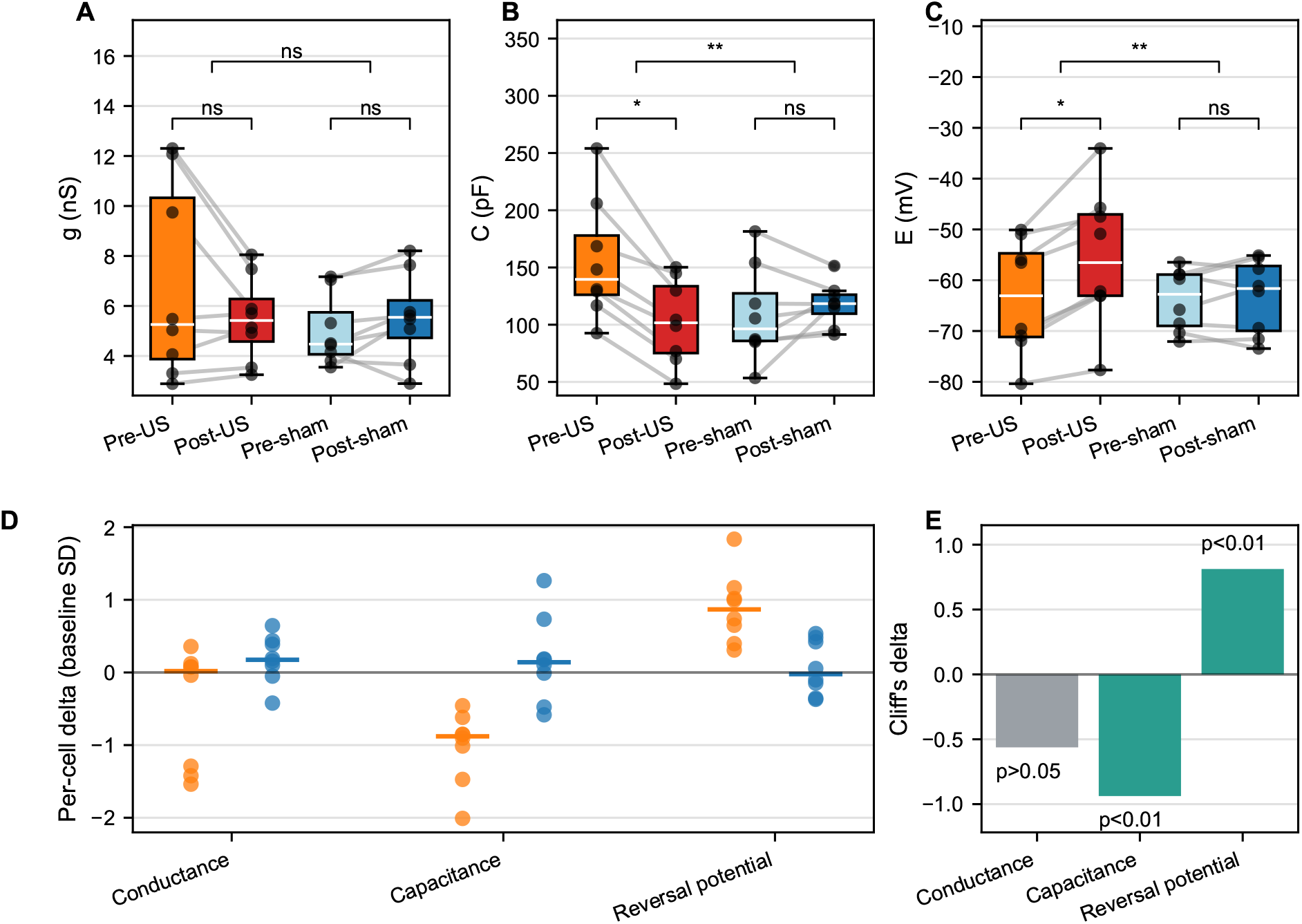
Effect of ultrasound on passive membrane parameters estimated from an RC fit to hyperpolarizing current steps. A–C. Input conductance *g*, membrane capacitance *C*, and resting (leak) reversal potential *E* before and after treatment in sonicated (orange) and sham (blue) cells; gray lines connect paired measurements within the same cell. D. Per-cell treatment deltas, each parameter divided by the standard deviation of its pre-treatment values across all cells so that quantities in different units share one axis (orange: ultrasound; blue: sham); horizontal bars mark group medians. E. Between-group effect sizes (Cliff’s *δ* ) for the treatment deltas, with FDR-adjusted exact permutation *p* values indicated as thresholds; exact values are given in the text. Within-group changes were tested with paired Wilcoxon signed-rank tests; between-group treatment effects with exact permutation tests on median deltas, Benjamini–Hochberg corrected across the three parameters.

**Figure 6.**
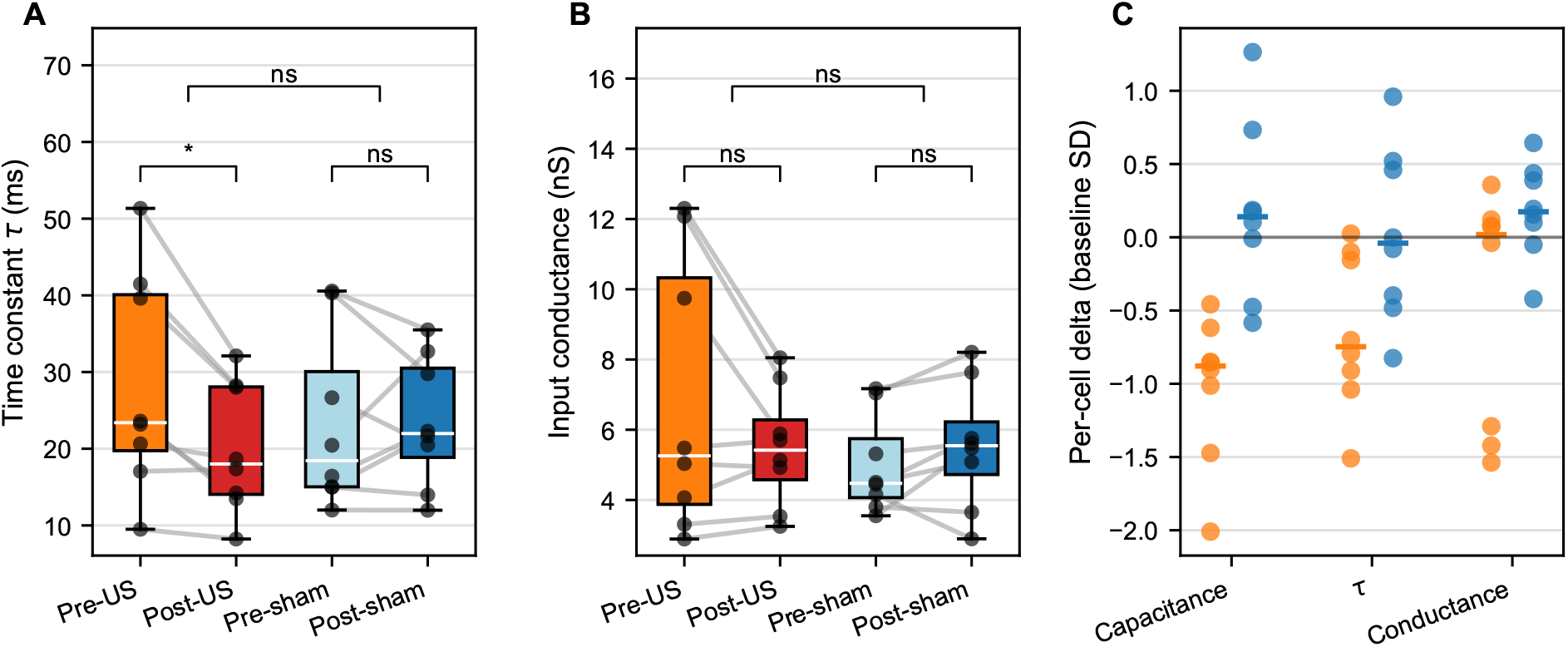
Per-cell membrane time constant and input conductance before and after treatment. A. Fitted membrane time constant 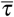. B. Input conductance *g*, shown alongside 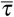 for direct comparison (the same values appear in Fig. 5A). C. Per-cell treatment deltas for capacitance, time constant and conductance, normalized as in Fig. 3E and Fig. 5D (orange: ultrasound; blue: sham); horizontal bars mark group medians. Because the capacitance is obtained as 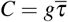 (see Methods), panel C shows that the capacitance change is carried by the time constant rather than by the conductance. Connected points in A and B are individual neurons (*n* = 8 per group) recorded before and after treatment; boxplots show the median, 25^th^ and 75^th^ percentiles, range, and outliers. Significance markers in A and B are FDR-corrected across the two parameters shown here (∗: *p*≤ 5 × 10^−2^; ns: not significant). Note the wide scatter of the sham changes in panel C, which includes one sham cell whose capacitance more than doubled.

## 4. Discussion

In summary, our *in vitro* LIFU experiments, performed in synaptically isolated primary mouse hippocampal neurons using current-clamp recordings, revealed a coherent set of persistent, cell-intrinsic changes measured approximately 20 min after a single 1 min exposure: a significant depolarization of the resting membrane potential, a significant decrease in spike amplitude, and a marked increase in membrane-potential noise, together with a pronounced reduction in membrane capacitance and a depolarizing shift in the leak reversal potential, while the input conductance remained unchanged. In the following we argue that these last two passive changes provide a parsimonious account of the others, and that the effect is intrinsic to the sonicated neuron.

A defining feature of our design is that glutamatergic and GABAergic transmission was pharmacologically blocked by applying CNQX, AP5, and bicuculline in the bath solution, so that the changes reported here are cell-intrinsic rather than network- or synapse-mediated. During initial testing without these blockers we observed large spontaneous bursts of action potentials that precluded reliable analysis (data not shown); their removal ensured that recorded activity reflected the intrinsic properties of the patched cell and the imposed current stimulation. This isolation also distinguishes our findings from the principal mechanisms proposed for the offline effects of ultrasound. Clennell et al. (2023) showed that the sustained increase in whole-cell potassium current they observed requires ionotropic glutamate receptor activation and is abolished by AMPA/NMDA receptor blockade; Watts et al. (2025) attributed lasting hippocampal changes to synaptic metaplasticity; and Kim et al. (2017) found the delayed rise in spontaneous spiking to depend on glutamate-receptor availability. Each of these routes is precluded in our preparation, yet robust offline changes persisted. Our blockade was also more stringent than the protocol that abolished the potassium-current effect. Clennell et al. pre-treated for 5 min with AP5 and NBQX and maintained antagonism during exposure and for 5 min afterwards, whereas our blockade was continuous across the whole experiment and additionally included a GABAergic antagonist. The triggers that remain available under our blockade are the receptor-independent mechanical and calcium pathways: activation of mechanosensitive channels and voltage-gated L-type calcium channels, which raise intracellular calcium and can themselves depolarize the membrane (Yoo et al., 2022; Fan et al., 2024). A receptor-independent, ultrasound-gated conductance of this kind has been measured directly in isolated crayfish axon under synaptic blockade (Lin et al., 2019). That conductance rose and decayed with the stimulus, whereas the change we report persists for at least 20 min after a 1 min exposure and is not accompanied by any rise in conductance.

Our results also refine the prevailing view that brief LIFU alters excitability without changing passive membrane properties. Clennell et al. (2021) reported enhanced excitability lasting up to 8 h in primary rat cortical neurons, accompanied by changes in action-potential kinetics but no detectable change in input resistance, membrane capacitance, time constant, or resting potential. In contrast, our within-cell paired design revealed consistent passive changes: reduced capacitance and a depolarized reversal potential in all eight sonicated cells, together with resting depolarization. We attribute this difference partly to statistical sensitivity: whereas Clennell et al. (2021) compared separately sonicated and sham populations (an unpaired design subject to the large between-cell variance typical of dissociated cultures), we re-patched the same neuron before and after sonication, accounting for inter-cell variability and exposing shifts that are uniform within cells yet modest relative to the population spread. Differences in capacitance estimation (a single hyperpolarizing step versus a multi-step exponential fit), in stimulation regime (850 kHz continuous wave versus 200 kHz pulsed), and in preparation (mouse hippocampal versus rat cortical) may also contribute. We therefore interpret the passive membrane change as a genuine intrinsic effect of LIFU rather than an inconsistency with earlier work.

Because capacitance estimates depend on how they are obtained, it is worth stating what our procedure measures. Golowasch et al. (2009) found that current-clamp step protocols, of the kind used here, recover total capacitance more faithfully than voltage-clamp step or ramp protocols, whose accuracy degrades as a neuron departs from isopotentiality. In a cell with dendrites, however, our estimate remains an effective rather than a total capacitance, for two reasons: a single exponential fitted over the first 50 ms is weighted toward the fast somatic relaxation and truncates slower dendritic components, and *C* is formed with the input conductance rather than with the amplitude coefficient of the slowest component, which is the divisor Golowasch et al. (2009) identify as the correct one. Both make our *C* a lower bound on total membrane capacitance. Gonzalez et al. (2025) quantified the size of this offset in dentate granule cells: replacing a monoexponential fit with a multiexponential fit and taking the slowest component raised the estimated capacitance in the same recordings from approximately 103 pF to approximately 133 pF.

A systematic offset of this kind does not by itself compromise a paired comparison, as shown in Gonzalez et al. (2025). Specifically, in dentate granule cells with genetically induced hypertrophy, two capacitance measures whose absolute values differed several-fold nonetheless reported statistically indistinguishable relative changes. Applying identical current steps, fitting window and estimator to the same neuron before and after treatment therefore removes the estimator-dependent part of the bias. However, if sonication altered the electrotonic structure of the cell, the distribution of time constants would also change, and the fraction of the slow relaxation captured within the 50 ms window used here need not be identical in the two recordings. The sham group bounds this concern, since those cells underwent the same transfer, re-patching and analysis without a significant change in capacitance, but it does not fully remove it.

The evidence for a LIFU-driven increase in firing was, by contrast, equivocal in our data. The post-treatment increase in the I–F slope reached significance within the ultrasound-treated cells, but the treatment-specific difference relative to sham was not statistically robust: it did not reach significance in our mixed-effects models, although the corresponding term in the repeated-measures ANOVA did. We therefore regard the increase in input–output gain as suggestive of, but not unequivocally attributable to, LIFU. One likely contributor is that the amplitude of our stimulating current steps did not extend beyond the range available in the shared analysis window, whereas Clennell et al. (2021) used currents up to 400 pA and observed significantly increased firing primarily above 250 pA; we cannot exclude that stronger between-group differences would emerge at higher current levels.

A central feature of our results is the concurrent reduction in membrane capacitance and depolarizing shift in the leak reversal potential, without a change in input conductance. Because the specific capacitance of neuronal membranes is nearly constant, at approximately 0.9 µFcm^−2^ (Gentet et al., 2000), a decrease of this size could in principle index a loss of effective membrane area. Our own measurements argue against that reading on its own. The membrane time constant, *τ* = *C/g* = *c*_m_*/g*_m_, is set by the specific capacitance *c*_m_ and the specific leak conductance *g*_m_, both per unit membrane area, and is therefore independent of that area. A loss of area at unchanged channel density lowers *C* and *g* in proportion and leaves *τ* untouched, yet the measured 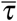 fell by about a third while *g* did not fall correspondingly (Fig. 6). At least one property of the membrane per unit area must therefore have changed: either the specific capacitance fell with the membrane area largely preserved, or the area fell and the specific leak conductance rose by roughly half, the two changes cancelling in *g*. Our data do not separate these. An increase in bilayer thickness would lower the specific capacitance directly, and has been proposed on independent grounds for ultrasound acting on crayfish motor axons and muscle fibres (Yu et al., 2023). The same reasoning applies to the leak reversal potential. *E* is the conductance-weighted mean of the reversal potentials of the pathways carrying the resting conductance, so it too is unchanged if every conductance scales with membrane area. The two quantities that shifted consistently across cells, 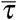 and *E*, are therefore both independent of membrane area, whereas *C* and *g* are not. A change in area alone would have moved *C* and *g* together and left 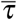 and *E* where they were. It also follows that the near-constant *g* does not imply an untouched resting conductance. The depolarized reversal requires its composition to have changed, while only the magnitude of the total conductance stayed the same. Membrane capacitance, long treated as a fixed quantity, is in fact dynamically regulated (varying up to two-fold across the day in hippocampal and cortical neurons (Severin et al., 2024)), so a persistent, sonication-induced reduction is biophysically plausible. We propose that both passive changes could arise from a single, lasting alteration of the membrane’s material state. Low-intensity ultrasound mechanically deforms the lipid bilayer, through intramembrane cavitation and bilayer-sonophore dynamics (Krasovitski et al., 2011; Plaksin et al., 2016) and directly measured membrane deflection (Vasan et al., 2022), and perturbs the cortical actin cytoskeleton to which the membrane is mechanically coupled (Mizrahi et al., 2012; Chen et al., 2014; Samandari et al., 2017). That a protein-free lipid bilayer can itself undergo persistent, conditioning-induced changes in area, thickness, and capacitance (Scott et al., 2022) shows that the membrane can retain such a change without protein machinery. A mechanically or calcium-triggered change in the thickness or lipid order of the bilayer would lower the specific capacitance directly; a reduction in membrane area, for example through calcium-dependent membrane retrieval (Wu et al., 2009), would lower capacitance only in combination with the rise in specific leak conductance noted above. In either case, an accompanying shift in the composition of the resting conductance toward a less K^+^-selective, more cation-permeable leak would depolarize the reversal potential. Importantly, because the same internal solution dialyzed the cytoplasm during both recordings, the equilibrium potentials of the major permeant ions were fixed; the depolarized reversal is therefore unlikely to reflect altered ion gradients and more plausibly a change in resting membrane permeability. The absence of any rise in input conductance argues against persistent membrane poration or damage, consistent with the preserved cellular morphology and firing. Sonoporation or seal injury would raise *g*, but it does not rise in any consistent way. The sham arm also bounds a further concern, that the second giga-seal rather than the sonication caused the change. Giga-seal formation is itself a mechanical stimulus to the neuron (Cepkenovic et al., 2023), but sham cells were transferred and re-patched under an identical protocol and showed no capacitance change (median Δ = +7.3 pF; Wilcoxon *p* = 0.461). We emphasize that this mechanism is a hypothesis motivated by our observations and the available literature, and that establishing it will require direct measurements of membrane area, thickness and permeability.

These two passive changes account for the remaining observations. The depolarizing shift in the leak reversal potential (≈+7.6 mV) closely matches the depolarization of the resting potential (≈ +5.9 mV), and the resulting steady depolarization would drive a fraction of voltage-gated sodium channels into steady-state inactivation, reducing action-potential amplitude (Bean, 2007). The reduced capacitance shortens the membrane time constant (*τ* = *C/g*) at unchanged conductance, speeding the approach to threshold, which predicts a shorter first-spike latency and a steeper input–output relation (in our data the latency change was not significant, and the gain increase was significant only within the sonicated group), and diminishes the low-pass filtering of membrane currents, which, together with any added stochastic leak, would increase membrane-potential noise (White et al., 2000; Severin et al., 2024).

Three features of our approach warrant emphasis. The synaptic isolation described above removed spontaneous network activity, and the within-cell repeated-measures design minimized the impact of inter-neuronal heterogeneity, allowing subtle intrinsic shifts to be detected with a modest sample. Because every measurement was made before or after the exposure and none during it, the recordings are also free by construction of the artifacts that contaminate intracellular electrophysiology performed while the transducer is driven (Collins and Mesce, 2020). A limitation of this design is that each neuron had to be patched twice; although we observed no gross loss of viability by light microscopy or basic electrophysiology, the reduced success rate on re-patching constrained the observation window to approximately 20 minutes. Our data cannot distinguish whether the capacitance reduction reflects a fall in specific capacitance, through a change in bilayer thickness or dielectric properties, or a loss of membrane area accompanied by a rise in specific leak conductance, nor identify the molecular basis of the reversal-potential shift. Additional studies with larger samples, longer follow-up intervals, and direct measurements of membrane area and permeability will be needed to test the proposed mechanism and to determine the duration and specificity of these effects.

## Additional information

### Data availability statement

All individual data points underlying the reported comparisons are shown in the figures. The original recordings and the analysis code are available from the corresponding author upon reasonable request.

### Author contributions

The experiments were performed at the Department of Physiology and Neurobiology, Eötvös Loránd University, Budapest, Hungary. M.S.: software, visualization, formal analysis, funding acquisition, writing – original draft; É.É.: investigation, data curation, formal analysis, writing – original draft; B.K.: formal analysis, investigation, writing – original draft; A.M.: formal analysis; V.H.: software, formal analysis; M.E.G.: cell culture preparation and maintenance; Z.S.: conceptualization, formal analysis, writing – review and editing; A.Sz.: methodology, data curation, investigation, writing – review and editing; T.K.: conceptualization, funding acquisition, supervision, writing – review and editing. All authors approved the final version of the manuscript; agree to be accountable for all aspects of the work in ensuring that questions related to the accuracy or integrity of any part of the work are appropriately investigated and resolved; and all persons designated as authors qualify for authorship, and all those who qualify for authorship are listed.

### Generative AI Statement

During the preparation of this work, the authors used GitHub Copilot (GitHub, Inc.; GPT-4.1) for coding assistance, Gemini (Google DeepMind; Gemini 3.1 Flash-Lite) for spell-checking the manuscript, and Claude (Claude Opus 5.5, Anthropic) for scientific-literature search, reference verification, and assistance with text editing, and with writing figure-generation code. After using these tools, the authors reviewed and edited the content as needed and take full responsibility for the content of the published work.

## Acknowledgements

We thank Barnabás Szűcs of Medi-CAD Kft (Miskolc, Hungary) for transducer manufacturing, and dr. Zsolt Molnár (Technical University, Budapest, Hungary) for the space scanner used for hydrophone measurements.

## Appendix

### A1.1. In-house-built ultrasound generator

Ultrasound stimulation for the sonication of cell cultures was performed using in-house-designed and in-house-built hardware (Fig. A1).

**Figure A1:**
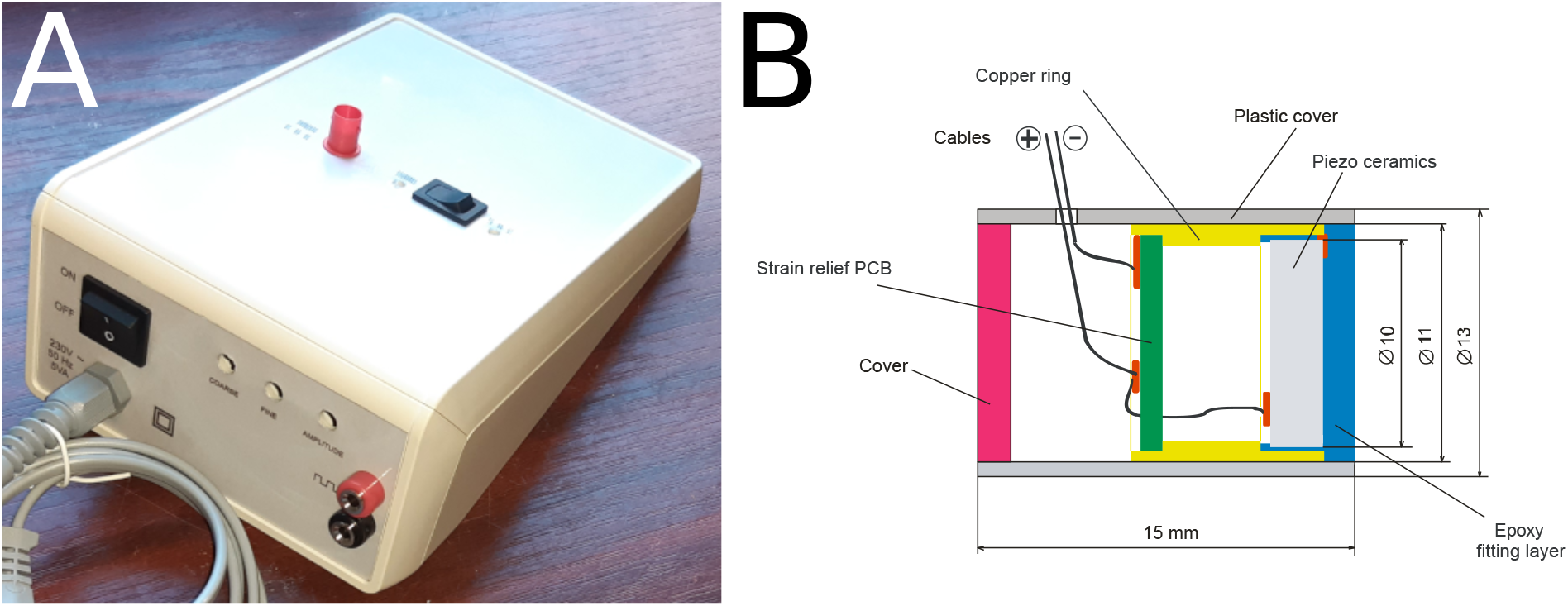
In-house-built ultrasound generator system. (A) Signal generator and amplifier capable of generating sinusoidal voltages in the 0.5–1.5 MHz frequency range with peak-to-peak amplitudes up to 40 V. (B) Schematic drawing of the ultrasound transducer designed to fit the small area of the coverslips.

